# Modulation of Type IV pili phenotypic plasticity through a novel Chaperone-Usher system in *Synechocystis sp.*

**DOI:** 10.1101/130278

**Authors:** Anchal Chandra, Lydia-Maria Joubert, Devaki Bhaya

**Affiliations:** Department of Plant Biology, Carnegie Institution for Science, 260 Panama Street, Stanford, CA940305, USA; CSIF, Stanford University Medical Center, Stanford, CA 94305-5301 USA

**Author notes:** Corresponding author: Name: Anchal Chandra Address: Carnegie Institute for Science 260 Panama Street, Stanford, CA-94305 and Name: Devaki Bhaya Address: Carnegie Institute for Science 260 Panama Street, Stanford, CA-94305.

**Keywords:** Phenotypic plasticity, Type-IV Pili, Chaperone-Usher, Phototaxis, *Synechocystis sp*., *PCC6803*, Social motility

## Abstract

Controlling the transition from a multicellular motile state to a sessile biofilm is an important eco-physiological decision for most prokaryotes, including cyanobacteria. Photosynthetic and bio geochemically significant cyanobacterium *Synechocystis sp.* PCC6803 (*Syn6803*) uses Type IV pili (TFP) for surface-associated motility and light-directed phototaxis. We report the identification of a novel Chaperone-Usher (CU) system in *Syn6803* that regulate secretion of minor pilins as a means of stabilizing TFP morphology. These secreted minor-pilins aid in modifying TFP morphology to suit the adhesion state by forming cell to surface contacts when motility is not required. This morphotype is structurally distinct from TFP assembled during motile phase. We further demonstrate by examining mutants lacking either the CU system or the minor-pilins, which produce aberrant TFP, that are morphologically and functionally distinct from wild-type (WT). Thus, here we report that in *Syn6803,* CU system work independent of TFP biogenesis machinery unlike reported for other pathogenic bacterial systems and contributes to provide multifunctional plasticity to TFP. cAMP levels play an important role in controlling this switch. This phenotypic plasticity exhibited by the TFP, in response to cAMP levels would allow cells and cellular communities to adapt to rapidly fluctuating environments by dynamically transitioning between motile and sessile states.

**Significance of this work:** How cyanobacterial communities cope with fluctuating or extreme environments is crucial in understanding their role in global carbon and nitrogen cycles. This work addresses the key question: how do cyanobacteria modulate external appendages, called Type IV pili, to effectively switch between motile and sessile biofilm states? We demonstrate that cells transition between forming strong cell-surface interactions indispensable for biofilm formation to forming cell-cell interactions that allow for coordinated movement crucial for social motility by functional/ structural modification of same TFP appendage. The second messenger, cAMP and a Chaperone-Usher secretion are indispensible to achieve these structural modifications of TFP and control the complex phenotypic transition. We have uncovered a strategy that *Syn*6803 has evolved to deal with molecular decision-making under uncertainty, which we call phenotypic plasticity. Here we demonstrate how a single motility appendage can be structurally modified to attain two antagonistic functions in order to meet the fluctuating environmental demands.

## Introduction

To sustain life in both terrestrial and aquatic environments, cyanobacteria can form multicellular light-driven motile communities or sessile, multi-species biofilms encased in an extracellular polysaccharide (EPS) matrix. In pathogenic bacteria, where the transition from a planktonic motile state to a sessile biofilm has been extensively studied and requires the expression of flagella, which is turned off as the community progresses into a biofilm state. Type IV pili (TFP) and EPS **^1^** are synthesized at this stage to facilitate surface adhesion instead of motility. There is an ordered progression towards a complex biofilm state**^2^** and the physiology of bacteria within a biofilm is distinct from the same cells in a planktonic state**^1^**. TFP in pathogenic bacteria are currently assigned a role only in the surface-associated adhesion required for biofilm related social behavior. Several proteins are required for the biogenesis, assembly and function of TFP**^3, 4^** which has been studied in the multicellular swarms of *Myxococcus. xanthus***^5^**, and in the pathogens *Pseudomonas aeruginosa***^6^**, *Neisseria gonorrhoeae*, *Escherichia coli* and *Vibrio cholerae***^7, 1^**.

In contrast, little is known about the regulation of social behavior of environmentally relevant phototrophs. Notably, many members in the phylum Cyanobacteria encode TFP**^8^** but none are reported to encode flagella**^9^**. This suggests that cyanobacteria are likely to use different strategies to switch between motile and sessile states. Here we report that the phenotypic plasticity of the TFP allows *Synechocystis* sp. PCC 6803 (hereafter *Syn6803*) cells to control switching between these states. *Syn6803* cells are capable of ‘twitching’ or ‘gliding motility’ over wet surfaces using TFP. *Syn6803* cells also exhibit motile social behavior that can be regulated by the direction, intensity and wavelength of light, a phenomenon known as phototaxis**^10^**. Mutants lacking structural and regulatory components of TFP exhibit a variety of motility phenotypes**^11, 12^**. For instance, *Syn6803* lacking *pilA1* (*sll1694*), which encodes pilin, the major structural component of TFP, are non-motile and lack TFP**^13^**. Phototaxis in *Syn6803* additionally requires multiple photoreceptors**^14, 15, 16, 17^** and a complex signaling network**^10, 17^.** Furthermore, cells that cannot synthesize adenylyl cyclase (Δ*cya1*) exhibit limited motility**^18, 19^**. In this work we demonstrate that the Δcya1 strain makes structurally aberrant TFP that might explain their limited motility and that addition of extracellular cAMP rescues TFP function and morphology, as well as motility.

In some bacteria, a dedicated secretion system comprised of a chaperone and usher (CU) protein is upregulated during biofilm formation. This system is required for the transport and assembly of secondary structures (fimbriae, curli**^20^**) at the cell surface that promote cell-adhesion. The CU pathway serves as a minimal secretion system that allows for protein secretion without energy (ATP) consumption**^21, 22, 23^**. We have identified a CU system in *Syn6803* that appears to play a novel role in assisting the modification of TFP rather than in the secretion of secondary adhesive appendages.

To identify the role of this novel CU system and function of its secreted products, we took several complementary approaches. We used biochemical and pull-down assays to show that the putative CU system was involved in secretion of minor-pilins and an adhesin-like protein to the cell surface which serve complementary, yet distinct roles, in modifying TFP and the cell surface. Finally, we exploited gene-specific mutants to help dissect the complex relationship between TFP function and motility phenotypes by using phototaxis assays and electron microscopy. We also determined that cAMP was involved in the regulation of motility via these secreted products.

Based on these results, we present a preliminary model to explain how cell communities transition between motile and sessile states. The model also highlights the central role of cAMP in regulating several key components, including a novel CU system and minor-pilins that modify the functional behavior of the TFP. We propose that *Syn*6803 can rapidly alter TFP function, which changes the propensity of pili to adhere to surfaces (or other cells) or to glide over surfaces. This phenotypic plasticity exhibited by the TFP allows cells and communities to deal with molecular decision-making under uncertainty.

## Results

### cAMP controls TFP assembly

cAMP is a ubiquitous second messenger involved in signal transduction in bacteria and eukaryotes. Adenylyl cyclase converts ATP to 3′, 5′-cyclic AMP (cAMP) which binds catabolite receptor protein (CRP). The cAMP-CRP complex in turn, activates or represses transcription of downstream genes by binding to a conserved palindromic motif (TGTGAN_6_TCACA)**^24^**. *Syn6803* encodes a CRP (Sycrp1; sll1371) and an adenylyl cyclase (Cya1; slr1991)**^19^**. When WT cells are spotted on soft agarose plates, motile cells phototax to the front of the drop and then move out making typical long finger-like projections. Δ*cya1* cells are motile but exhibit an aberrant behavior; they aggregate at the front of the drop but rarely move beyond the front (**Fig. 1A**, **upper panel**). This impaired motility phenotype can be rescued in Δ*cya1* cells by the addition of extracellular cAMP (0.1mM) (**Fig. 1A**, **lower panel**). To investigate the cause of this impaired motility, we first examined the TFP of WT and Δ*cya1* cells by negative staining followed by transmission electron microscopy (TEM) (**Fig. 1B**, **upper right panel**). WT cells (**Fig. 1B**, **upper left panel**) typically exhibited several long TFP (> 2μm, 0.006± 0.002mm width), which were often connected to neighboring cells. Surprisingly, Δ*cya1* cells had none of these long TFP but only a few (∼4-5) very short, tapered (∼0.2μm length, 0.03 mm± 0.013 width) surface structures.

**Fig.1:**
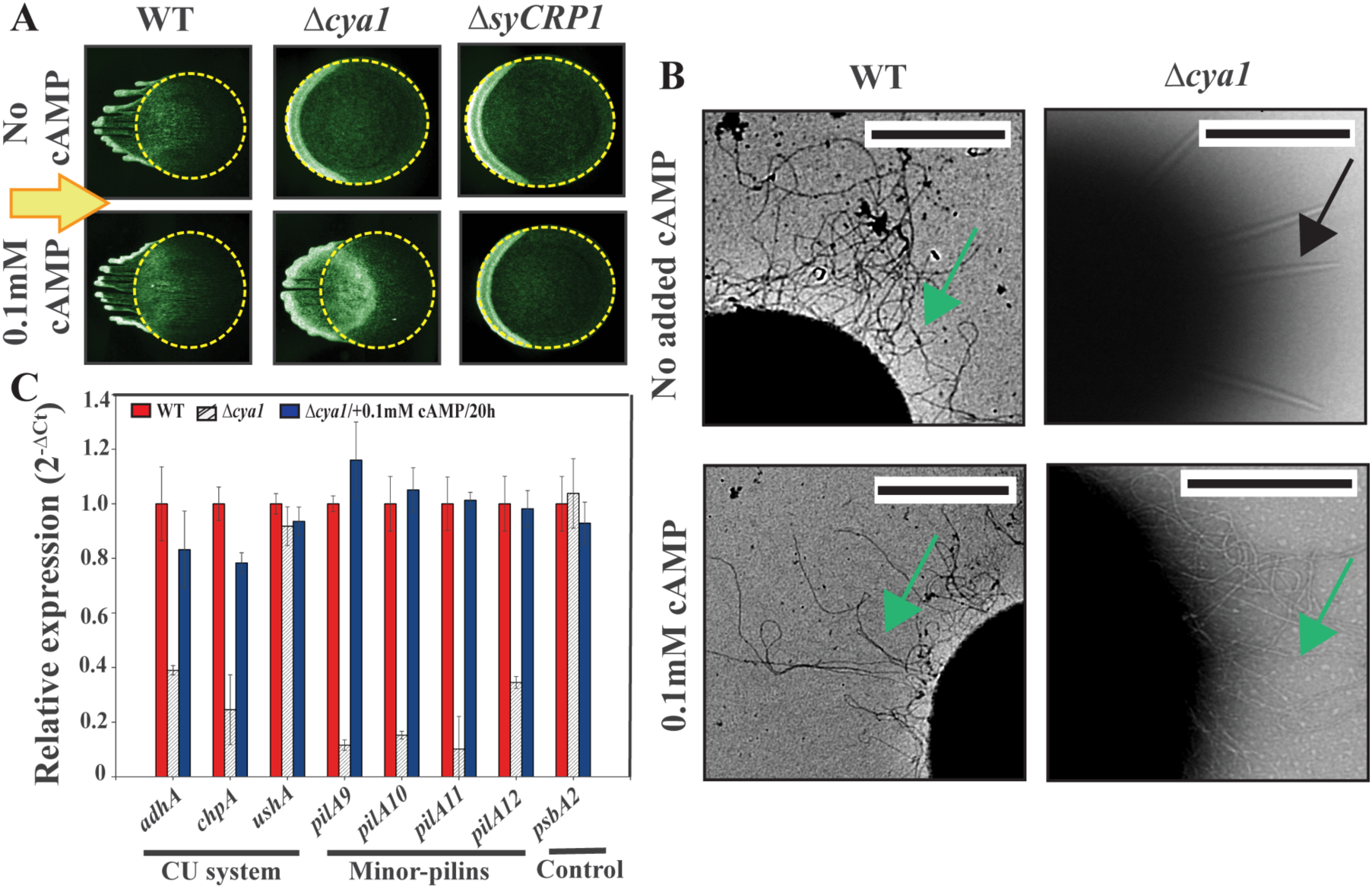
Effect of cAMP on Phototaxis, TFP morphology and transcription in WT and Δ *cya1* and *Δ sycrp1* (complementation assays). (**A**) *Effect of cAMP on phototaxis*. **Left panel:** WT cells, **Middle panel:** Δ*cya1* cells, **Right panel:** Δ*syCRP1* cells. Motility phenotype after 48h in unidirectional white light 30 μmol m–2 s–1 on the 0.4% agarose plates in the absence (**upper panels**) or presence (**lower panels**) of 0.1mM cAMP. Initial outline of drop (diameter 3-4mm) highlighted in yellow. WT cells make typical finger like projections of groups of moving cells. In the absence of cAMP, both ΔsyCRP1 and ΔCya1 cells accumulate (seen as darker crescent) at front of drop, In the presence of cAMP, Δcya1 cells regain the ability to form finger-like projections similar to WT. (**B**) *Effect of cAMP on TFP morphology*: Electron micrograph of negatively stained WT (**left**) Δ*cya1* cell (**right**) in the absence (**upper**) or presence 0.1mM cAMP for 20h (**lower**). Stubby, short TFP (black arrow); typical Type IV pili produced in WT and after cAMP addition (green arrow). Scale bar=1μm. (**C**) *Effect of cAMP on transcript levels*: Bar graph of relative mRNA transcript levels of *chpA*, *adhA*, and minor-pilin genes in WT cells (red), Δcya1 cells (striped) and Δcya1 cells after incubation with 0.1mM cAMP (blue) quantified by real time PCR (n=3 independent experiments). Error bars denote mean ± s.d where specified. *psbA2* serves as a reference control. C_t_ values of the reference control were used to normalize the C_ts_ for the gene of interest (ΔΔC_t_).

We reasoned that if cAMP was able to rescue the motility behavior of the *Δcya1* cells we should also be able to observe the short structures transition back to long TFP. First, we verified that cAMP could enter cells when added extracellularly, by using a competitive enzyme immunoassay (**Fig. S1a**). This showed that there was a slow increase in intracellular cAMP levels over time, after addition of cAMP to the agarose plate. When we supplemented cells with extracellular cAMP (0.1mM) for 20 hours (**Fig. 1B**, **lower panels**), and examined TEM images, we found that cells now exhibited TFP similar to WT cells. This indicated that cAMP actively controls the assembly of long TFP and in its absence short, but functional, TFP are formed. Thus, cAMP might mediate correct TFP assembly either by directly controlling pilin expression levels or by controlling the synthesis of additional proteins that are required for TFP function or stability. To distinguish between these two possibilities, we used qRT-PCR to check if *pilA1* (encoding the major TFP pilin) and other previously identified pilin-like proteins (*pilA2-A8*) were controlled by cAMP. There was no change in *pilA1* levels or other pilin assembly genes in Δ*cya1* cells, supplemented with cAMP (**Fig. S1b**), indicating that cAMP does not directly regulate pilin (PilA1) synthesis. Alternatively, cAMP could regulate TFP assembly by controlling the activity of other genes. To identify these putative genes, we used two approaches. First, we identified genes and operons that contained predicted upstream conserved CRP binding sites**^23^**. Second, we focused on genes that had been reported to be strongly down-regulated in Δ*Cya1* cells**^25 26^**. Two putative operons met these criteria, the first contained two genes (slr1667, slr1668) and the other contained four genes (slr2015, slr2016, slr2017 and slr2018). Using qRT-PCR, we showed that addition of extracellular cAMP restored the expression levels of all of these genes, close to WT levels in *ΔCya1* cells (**Fig. 1C**). This suggested that some or all six of these gene products might affect TFP-dependent motility and so we used complementary approaches to identify their specific roles.

### Identification of a novel Chaperone-Usher (CU) system in *Syn6803* that is regulated by cAMP

The genes slr1667 and slr1668 comprise an operon with a canonical cAMP receptor (SyCRP1) binding site (**<u>TGTGA</u>**TCTGGG**<u>TCACA</u>**) upstream (**Fig. 2A**). slr1668 is highly homologous to the chaperone protein of the CU system**^21, 27, 28^** and shows 87% overall conservation and greater than 67% conservation in the structural motifs and residues characteristic of the FGS sub-family members (chaperone proteins that contain a **s**horter loop between **F**1 and **G**1 beta strands) (**Fig. S2a**) that is vital for chaperone function. An overlay of the crystal structure of the evolutionarily related PapD, FimC chaperones with a homology model of slr1668 revealed two immunoglobulin-like domains characteristic of the chaperone family proteins (**Fig. S2b**). slr1667 encodes a protein of 178aa with very low homology to a fimbrial protein (**Fig. S2d)**, although the N-terminal contains a 25aa signal peptide and a typical Sec-like secretion signature motif. Using a Hidden Markov model (HMM) we identified divergent repeat units (24-40aa and 47-64aa) (**Fig. S2c**). Such repeats are characteristic of fimbrial tip adhesins secreted via the CU pathway and suggest that slr1667 may be secreted via the CU system. Previously, it was shown by immunocytochemical analysis that slr1667 is localized at the cell surface, while slr1668 is localized in the cell periplasm**^29^**. Therefore, we refer to slr1668 as ‘*chpA’* (chaperone) and slr1667 as ‘*adhA’* (adhesin).

**Fig. 2:**
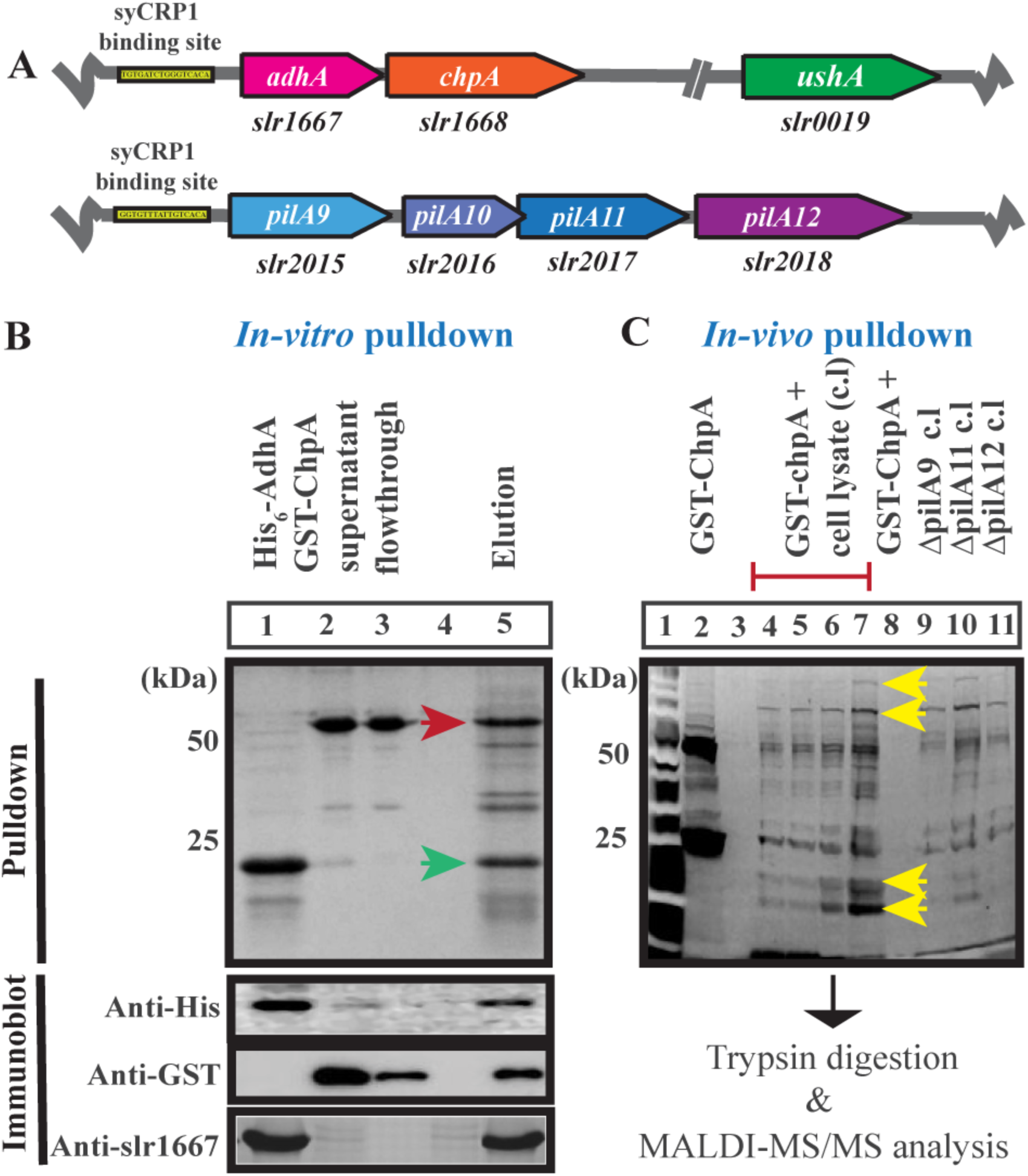
Gene loci regulated by cAMP/Sycrp1 and protein-protein interaction assays. (**A**) *Organization of gene loci regulated by cAMP* **Top row:** CU system genes **Bottom row:** minor-pilins genes (*pilA9*, *pilA10*, *pilA11*, *pilA12*). SyCRP1 binding site is indicated in yellow box. (**B**) *In-vitro protein interaction* analyzed by SDS-PAGE. Upper panel: Lane (1) Purified His_6_-AdhA, (2) GST-ChpA loading, (3) flowthrough, (5) co-eluted His_6_-AdhA (green arrow∼25kDa) and GST-ChpA (red arrow∼50kDa). **Lower panel:** Immunoblot with anti-His, anti-GST and anti-AdhA antiserum. (**C**) *In-vivo GST pulldown assay.* **Upper panel:** Lane (1) protein ladder (2) Glutathione beads with purified GST-ChpA, lane (L4-7) Elution from samples containing beads with GST-ChpA and whole cell lysate of WT *Syn6803* cells. Each of four lanes contains increasing volume of eluted samples. Lanes containing eluted samples of GST-ChpA loaded with whole cell lysates of Δ*pilA9* (L9), Δ*pilA10* (L10), Δ*pilA12* (L11) strains. Yellow arrows indicate samples analyzed using MALDI-MS/MS.

Typically, genes encoding CU components are organized in an operon that includes three proteins: the chaperone, the usher and a fimbrial subunit, such as an adhesin **^21^**. In *Syn6803,* we identified a putative usher homolog, slr0019 which had 30% identity to the usher, FimD (**Fig. S2e**) but it is not adjacent to ‘*adhA’* and ‘*chpA’*. However, in other sequenced cyanobacterial genomes, the three proteins, chaperone, usher and adhesin are in predicted operons (**Fig. S2f**). We thus refer slr0019 as the usher or ‘*ushA’*. Notably, all the cyanobacterial genomes encoding a CU system also contained homologues of pilin biosynthesis genes (**Fig. S2g**) suggesting that they may have inter-related functions. Thus, *Syn*6803 have a putative CU system consisting minimally of ChpA, AdhA and UshA. Next we investigated the role of this putative CU system in motility.

### The Chaperone-Usher system secretes AdhA and minor-pilins (slr2015-slr2018)

To investigate if the chaperone, ChpA, was involved in transport of the putative adhesin, AdhA, we performed *in-vitro* pulldown assays using His tagged-AdhA and GST-ChpA. ChpA forms a stable complex with AdhA (**Fig. 2B**) suggesting that AdhA is transported out of the cell by the CU system, which is also consistent with its localization to the cell-surface**^29^**. Next, to establish if proteins other than AdhA were transported by the CU pathway, we performed *in-vivo* pulldowns (**Fig. 2C**). *Syn6803* whole cell lysate was incubated with GST-ChpA immobilized on GST magnetic beads, as bait. Several proteins were pulled down by GST-ChpA. Protein bands that were clearly visible with Sypro-Ruby stain were excised from SDS-PAGE and processed for analysis by mass mapping (MALDI-MS/MS). This clearly identified slr2018 as an interacting partner of ChpA (**Fig. 2C**). Low levels of other proteins could be seen in the ChpA pulldown assay but were hard to identify with MALDI-MS/MS (based on their size, these may have included previously identified slr2015, slr2017). The slr2018 polypeptide (∼96kDa) is encoded by the last gene in a putative operon that includes slr2015, slr2016 and slr2017. As described earlier, slr2018 is in a putative operon which is likely to be controlled by cAMP based on genetic and bioinformatics.

The N-terminal hydrophobic regions of slr2015, slr2016, slr2017 exhibit significant conservation to Type IV prepilins from Gram-negative pathogens and cyanobacteria (**Fig. S2h**) with a conserved peptidase cleavage site (**G**_F/V/Y_X<u>L</u>X**E**) recognized by the PilD signal peptidase**^3^**. slr2018 gene product has a somewhat unconventional N-terminal region (**G**_F_XXX**E**). These gene products were previously identified as PilA9 (slr2015), PilA10 (slr2016), PilA11 (slr2017)**^11^** and we extended the same nomenclature to include slr2018 as ‘PilA12’. Based on this preliminary identification, we refer to these as ‘minor-pilins’. Minor-pilins have been identified for other pathogenic bacterial systems and play an essential role in priming TFP assembly in *P. aeruginosa* **^30^**.

### Mutants in the CU system and minor-pilins display different motility phenotypes

To further probe the interactions between the CU system and the minor pilins we examined the motility behavior of the *ΔushA, ΔchpA* and*Δ adhA* mutants. We also took advantage of existing gene-specific, nonpolar mutants of minor-pilins, *pilA9*, *pilA11* and *pilA12***^13, 26^** to further identify the exact role of minor-pilins in *Syn6803* in TFP mediated motility.

#### ΔushA cells exhibit hyper-motile behavior

We reasoned that if the CU pathway and its partner proteins were critical for motility, mutants in this pathway should exhibit motility defects. We characterized the motility phenotypes of *ΔchpA*, *ΔadhA* and *ΔushA* mutants and the minor-pilin mutants, slr2015 (*pilA9*), slr2017 (*pilA11*) and slr2018 (*pilA12*) that were generated either by double homologous recombination or by transposon mutagenesis (**Table** S**1**, S**2**). Δ*ushA* cells exhibited two unique motility behaviors (**Fig. 3A**). As expected, WT cells phototaxed to the front of the drop and then after ∼24-30h, emerged as finger-like projections of motile cells. Surprisingly, Δ*ushA* cells, developed finger-like projections much faster than the WT cells, as early as 15h (**Fig. 3A, left panel, Fig. S3a**). This was consistent with the quantification of the mean velocity and speed of the WT and Δ*ushA* cells, over time. For instance, at 12h, the velocity of Δ*ushA* cells was thrice that of WT cells (**Fig. 3B, 3C, Fig. S3b, Table S3**). Second, we routinely observed that although the majority of motile WT cells phototaxed to the drop front, many small groups of cells remained “stuck” on the agarose. However, the Δ*ushA* mutants did not appear to “stick” to the surface (**Fig. 3A, right panel**). This suggested that there might be cell-surface protein(s) that makes the cell “sticky” in WT cells, which are absent from the cell surface of Δ*ushA* cells. Consequently, cells do not adhere to other cells or to the agarose surface, resulting in a ‘hyper-motile’ phenotype.

**Fig. 3:**
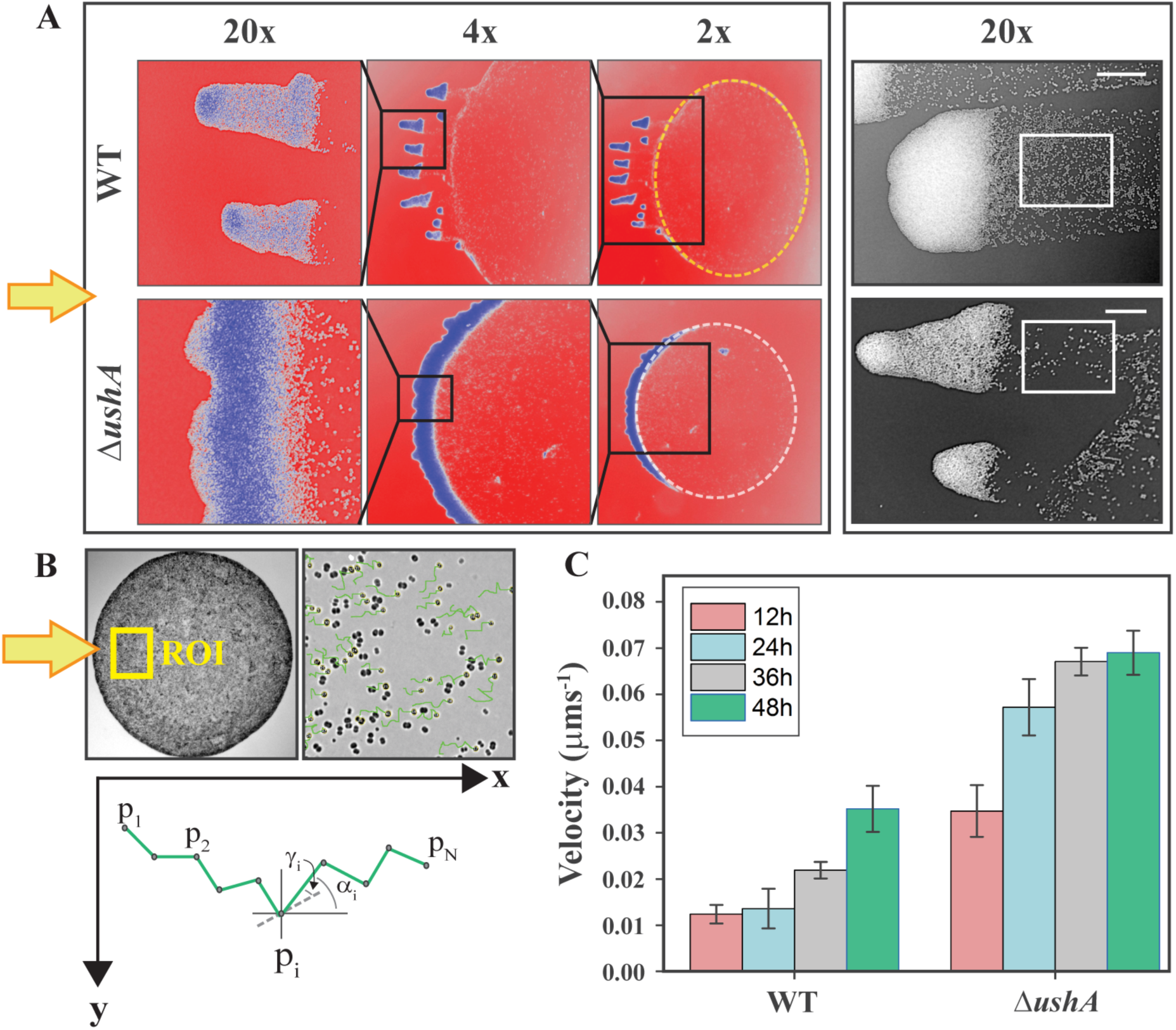
Hyper-motile Δ *ushA* strain display different motility phenotype under unidirectional light. (**A**) *Agarose motility assays*: **Left panel:** WT (upper row) and Δ*ushA* (lower row) after 20h motility assay (Arrow indicates direction of light). A small region (black box) from the drop front is magnified at 4x (middle) and 20x (left) to show relative density and movement of cells after first 20h of white light exposure. **Right panel:** 20X magnification of finger front after 48h depicting enhanced collective movement in Δ*ushA* cells (lower) versus WT (upper). See white box. Note how almost no cells are remain in the Δ*ushA* compared to WT. Scale bar∼ 20μm. (**B**) **Upper:** Example phase contrast image of the region of the drop used for quantification of average speed and velocity. Scale bar=20μm. **Lower:** The drawing shows a sample trajectory consisting of N points p_i_= (x_i_; y_i_). (**C**) Quantification of mean velocity (in WT versus Δ*ushA)* during different time points (12-48h). Each bar graph represents measurements of 40-50 individual cells for each strain at every time point.

#### Minor-pilin mutants are non-motile

In contrast to the hyper-motile Δ*ushA* cells, the three minor-pilin mutants (Δ*pilA9*, Δ*pilA11*, Δ*pilA12*) were completely non-motile (**Fig. S3c**), based both on the phototaxis assay and by measurements of cell movement that indicated that there was no displacement over time. These striking differences in the motility behavior of the mutants, ranging from hyper-motile *ushA* cells to non-motile minor pilin mutants, are apparently contradictory, particularly if they are in the same pathway. To reconcile the different phenotypes exhibited by the mutants, we examined other aspects such as (i) TFP biosynthesis, (ii) the morphology and assembly of the TFP and (iii) the ability of TFP to make cell-cell contacts.

### TFP synthesis, morphology and activity in CU and minor-pilin mutants

#### (i) *TFP synthesis is <u>not</u> affected in the CU and minor-pilin mutants*

To investigate whether the TFP biogenesis machinery or function depends on the CU pathway and/or on the minor-pilins, we analyzed the expression levels of several *pil* genes in the Δ*ushA* and minor-pilin mutants (Δ*pilA9*, Δ*pilA11* and Δ*pilA12*) via qRT-PCR. A core set of conserved proteins required for pilus biosynthesis including PilM, PilN, PilO, PilQ (pilus assembly), PilT PilB1(motor proteins), PilA1, PilA2 (pilins) are found in *Syn6803* and their role in pilus assembly has also been inferred from mutant phenotypes**^3, 11, 12^**. We measured mRNA levels of the following gene homologues: slr1277 (*pilQ*), slr1274 (*pilM*), slr1275 (*pilN*), slr1276 (*pilO*), slr0063 (*pilB1*), sll1694 (*pilA1*), sll1695 (*pilA2*), in the WT and in the mutants. We found that mRNA levels of pilins and components of pilus biogenesis in the mutants were comparable to WT levels (**Fig. S1b, c**). Thus, pilus biosynthesis appears to be unaffected in the CU and minor-pilin mutants and cannot explain the various motility phenotypes.

#### *(ii) TFP morphology and assembly is strongly affected in the* Δ*ushA* and minor pilin *mutant strains*

Next, we used high resolution TEM to investigate if the distinct motility defects in the mutants could be correlated with changes in TFP morphology (**Fig. 4A**). WT cells have several long TFP (Avg. number 15-20/cell, Avg. length = 2.5μm ± 0.05, Avg. width = 0.007μm± 0.002). The hypermotile Δ*ushA* cells completely lacked long TFP but short stubby structures (Avg. length= 0.7mm ± 0.2; Avg. width = 0.02 mm± 0.005) with tapered ends were consistently observed. These TFP were morphologically similar to those previously observed in Δ*cya1* cells (**Fig. 1B**). On average, there were 5-8 evenly distributed short stubby structures per cell, which are apparently functional, since Δ*ushA* is hyper-motile. Interestingly, the non-motile minor-pilin mutants also exhibited these short, stubby TFP (Avg. length= 0.78mm± 0.21 and 0.63mm ± 0.2). (**Fig. 4B, lower bar graphs)** shows the distribution of TFP and cell sizes (**Fig. S4a)**. This clearly indicated that some of the differences in TFP function and motility might be due to different TFP morpho-types (long TFP versus short, stubby TFP). This in turn, might affect their ability to form TFP-dependent intercellular cell-cell contacts.

**Fig. 4:**
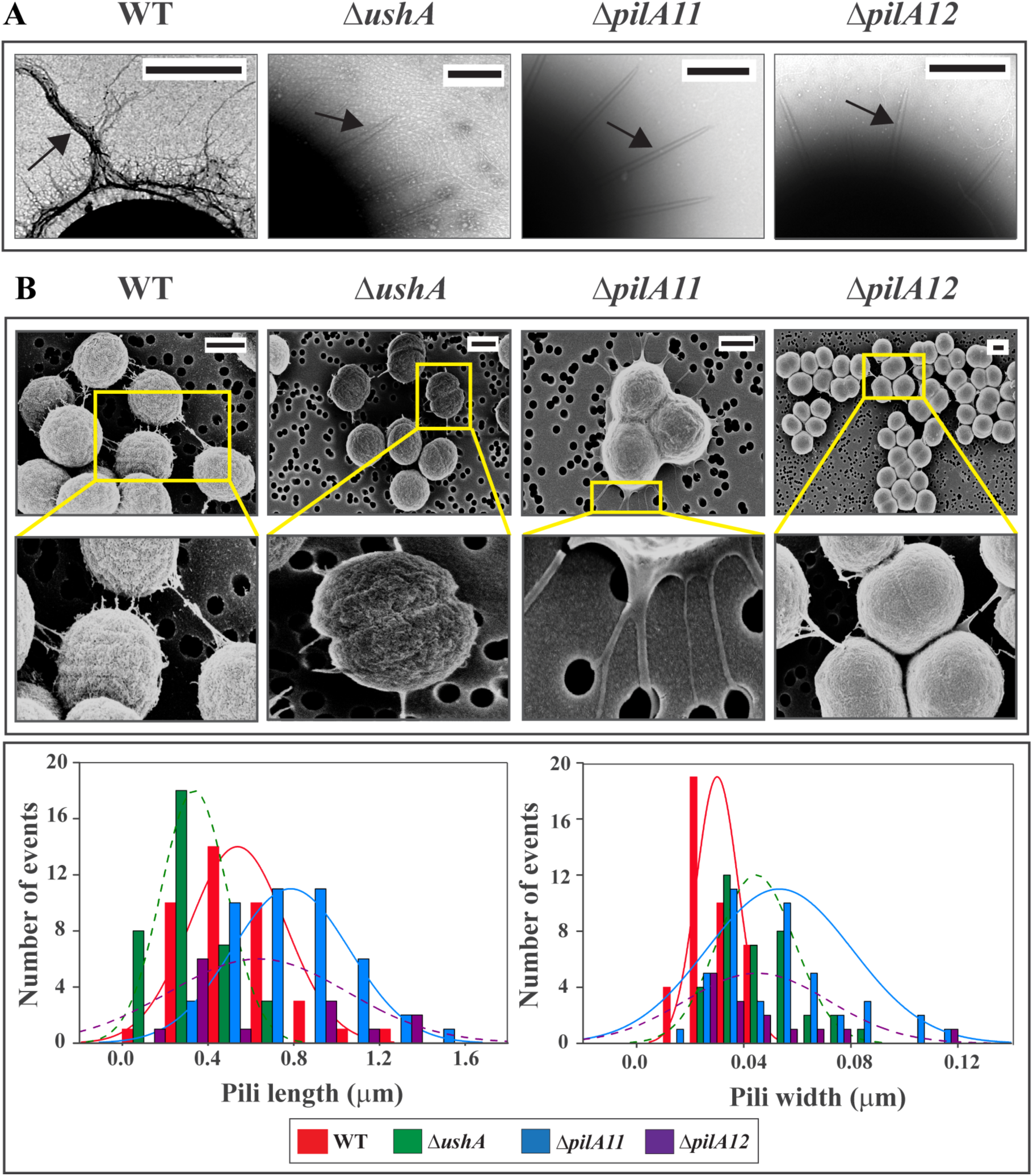
Identification of two TFP morphotypes in WT and mutants. (**A**) *Electron micrographs of negatively stained cells*: WT, Δ*ushA*, *ΔpilA11* and *ΔpilA12* after exposure to unidirectional light on motility plates. Scale 1 μm. Black arrows indicate representative normal TFP. (**B**) *Scanning electron microscopy* **Upper:** SEMs images showing cell-cell interaction (WT, Δ*ushA*) and cell-substrate interaction (Δ*pilA11* and Δ*pilA12*). Respective magnified images (yellow boxes) are shown below to WT and mutants. **Lower:** Left graph: Pilus length distribution in WT and mutant strains, Right graph: Pili width distribution.

#### (iii) Mutants exhibit an alteration in the ability to form cell-cell contacts via TFP in the mutants

We then used field emission scanning electron micrography (FE-SEM) to examine TFP morphology and cell-cell surfaces, which revealed another facet of TFP functionality. In WT and Δ*ushA* strains, TFP appear to form strong “cell to cell contacts” which originated around the cell body (**Fig. 4B**). In the minor-pilin mutants, the short TFP appeared to be mostly present at the base of the cell and were involved in forming strong contact with the grid surface rather than cell-surface. The cells were associated in large clusters (**Fig. 4B**) and had a much smoother cell surface than WT (**Fig. S4b**).

Consistent with this observation, the crystal violet absorbance assay, a semi-quantitative way to assess cell adherence or biofilm formation, showed that the minor-pilin mutants all had significant adhesion to glass culture vessels, relative to WT cells. In contrast, the usher (Δ*ushA*), chaperone (Δ*chpA*) and Δ*adhA* showed greatly reduced (10-30 fold) adherence (**Fig. S4c**). This suggests that secretion of AdhA via CU system is crucial for adhesion of cells. Conversely, in the absence of AdhA, cells experience no counter-force against directed motility and hence are hyper-motile (Δ*ushA*).

## Discussion

### A new role for a chaperone usher system in TFP assembly

We uncovered a novel role of CU (Chaperone-Usher) system in *Syn6803* which modifies TFP function rather than being involved in secondary filament synthesis as described for most pathogenic bacteria. The *Syn6803* CU system secretes minor-pilins that alter TFP morphology and hence their functionality. In the absence of the usher protein, short TFP are produced but the cells are as motile as WT cells suggesting that long TFP are not strictly necessary for surface motility. In support of this, it has been shown that *P.aeruginosa* mutants with very short pili (that are incapable of being sheared) are motile^31^. The CU system also transports AdhA, a cell adhesion/cell surface protein that serves as a crucial component in controlling reversibility of TFP function such that cells can transition between a motile and a sessile state. The ΔushA mutant does not transport the putative adhesin (AdhA), to the cell surface. This results in cells that do not appear to stick to the surface (Fig. S4c) and are hyper-motile relative to WT cells. Such a CU pathway would allow cells to modify TFP function in an energy-independent pathway as well as the cell surface properties in response to cAMP levels. This could allow cyanobacterial populations to make rapid and reversible decisions in response to unpredictable fluctuating environmental variables such as light and nutrients.

### Switching between pilus morphotypes allows for motile or sedentary lifestyles

Like other bacteria, *Syn6803* undergoes a transition between a motile and sessile (biofilm) state. We demonstrate that TFP in *Syn6803* upon undergoing modification via minor-pilins represent two different morphotypes-long TFP can form strong ‘cell to cell contacts’ (Fig. 4B, WT and ΔushA) thus facilitate social motility. Second, are the short TFP (Fig. 4B, ΔpilA11, ΔpilA12; FESEM images) that can form strong cell-surface interactions thus facilitating a stable, irreversible transition from motile to sedentary state. This is corroborated by the fact that all the minor-pilin mutants which have short pili appear to attach much more strongly to glass surfaces than WT cells (Fig. S4c). The presence of long TFP might physically obstruct contact with the surface, which can be achieved more effectively by short pili^32^. An alternative explanation is that frequent extension/retraction of dynamic long TFP; keep the cells in motion hence preventing the cells from achieving efficient surface contact^32, 33^. This scenario is in agreement with the finding that surface attachment properties of longer versus shorter type-I and TFP in *Xylella fastidiosa* are distinct^33^.

### Are motility and adhesion mutually exclusive processes?

Surface motility is a complex trait that involves a dynamic interplay between adhesiveness (irreversible attachment to surface) and TFP-driven directional movement in presence of light. The net displacement of cells is an outcome of these two antagonistic forces; i.e. reversible adhesion versus directed movement ^32, 34^. The apparent inconsistency between the motility behavior, TFP morphologies and surface adhesion abilities of the various mutants here provides a logical explanation for the distinct yet collaborative role of minor-pilins and AdhA proteins in mediating two processes-motility and adhesion. Motile WT cells assemble long functional TFP and have sufficient AdhA expression on the cell surface. On the other hand, non-motile minor-pilin mutants express same levels of AdhA on the cell surface as WT thus making them highly adherent. However, they lack the ability to secrete minor-pilins that modify/stabilize assembly of long TFP resulting in the short pilus phenotype. Conversely, the surface of hyper-motile ΔushA cells and Δcya1 mutants lack AdhA and minor-pilins which results in short pili and reduced surface adhesion ability; both these phenotypes allow them to move faster than WT cells. The crystal violet binding assay also corroborated that WT cells and minor pilins have stronger cell-surface adhesiveness compared to the ΔushA cells. This leads to the conclusion that short TFP pili are sufficient for motility (Fig. 3A, B), but the ability to adhere to the surface is controlled by AdhA levels. We propose that AdhA that is secreted via the usher pathway to the cell surface^28^ and exhibits similarity to fimbrial tip adhesins is crucial for cells to make this decision during the transition stage.

This scenario is in contrast to what has been proposed in most other bacterial systems, where motility and adhesion are antagonistic states and are inversely coordinated^35, 36, 37, 38^, In *Syn6803*, we propose there is a dynamic balance between motility and surface adhesion which is required for optimal movement over wet surfaces. We suggest that because surface-dependent motility (here phototaxis) is a group or social behavior, the dynamics cannot be fully explained simply based on the behavior of individual cells. Motility is enhanced by interacting with neighboring cells, mediated by long retractile TFP that can enhance attachment between cells and this enhances certain social behaviors^35^. The dynamic nature of these cellular interactions as seen in motility movies^17^ also support the idea that cells need to be “sticky” and “motile” and is maintained by minor pilins and AdhA levels. Finally, this delicate balance is controlled by cAMP levels. (see Model).

### A model of phenotypic plasticity in Syn6803

Based on our results, we present a model to explain how *Syn6803* dynamically switches between motile and sessile states. cAMP acts as a master regulator to control expression of minor-pilins (PilA9-PilA12) and the CU system (minimally comprised of an usher (UshA), chaperone (ChpA) and adhesion substance (AdhA). Secretion of minor pilins and AdhA to the outside of cell requires CU pathway (**Fig. 5A-upper**). Minor pilins secreted via the CU system help to stabilize TFP and ensure assembly of long TFP that form effective contacts between neighboring cells that is quite crucial for assisted group motility.

**Fig. 5:**
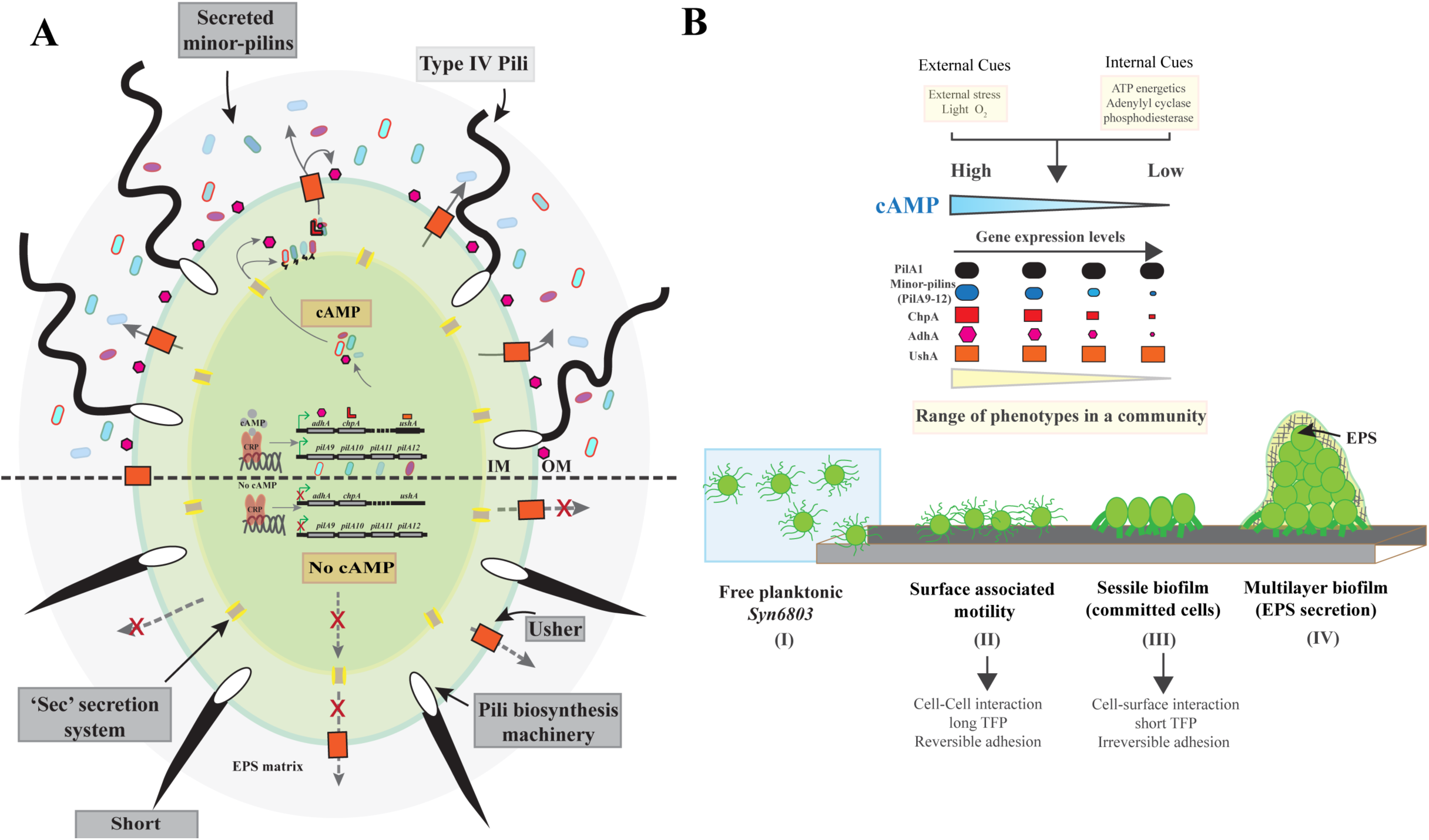
Model of cAMP regulation of TFP phenotypic plasticity. (**A**) Individual cell (**B**) Communities showing how dynamic range of effects can be created.

Transition from surface-associated motility to forming sessile biofilms involves two crucial steps (**Fig. 5B**). The first step involves cells making initial but potentially reversible contacts with surface. Before this stage, cells are still capable of switching back to the motile phase, based on environmental conditions. AdhA expression on the cell surface governs this decision of reversible attachment of cells. The second step of biofilm progression involves those cells that are committed to forming stationary biofilms and form irreversible cell-surface contacts. In the absence or low concentrations of cAMP (**Fig. 5A-lower**), minor-pilins and AdhA are either or absent or expressed at low levels for cells to form short stubby TFP. These functionally modified TFPs are enough to form strong cell to surface contacts to ensure irreversibility and hence commitment to biofilm formation. Future experiments to test these hypotheses, particularly how cell communities control the transition between the motile lifestyle and sessile biofilms will further substantiate our claims.

## Material and methods

### Growth and culture conditions

Freshwater *Synechocystis sp*. PCC 6803 (*Syn6803*) strain (Berkeley) was maintained in BG-11 medium supplemented with 5mM TES and 10mM NaHCO3 at 30° C with continuous shaking at 100rpm under aerobic conditions and warm white lights (25-30 μmol photons/ m^-1^ s^-1^). All experiments were performed with cells in exponential phase (O.D_730_= 0.6-0.8). For cell adhesion assays, cells were cultured on bench top without shaking under similar growth conditions.

### Transcription analysis by quantitative Real-time RT-PCR (qRT-PCR)

RNA extraction, cDNA synthesis and qRT-PCR measurements were performed as described here**^49^**.

### Agarose motility assays

Motility assays under directed unidirectional light were carried out at 30°C on 0.4% (w/v) Difco agar in BG-11 in 50mm plastic petri dishes (BD falcon-25369022). One microliter of cells (OD_730_ = 0.6-0.8) was placed in the center of a plate. A white LED was used as the light source. The incident light intensity as measured by LI-COR light meter was set to approximately 25μmol photons/ m-1 s-1. For time-lapse video measurements and single-cell tracking analysis see **SI Materials and methods**, Time-lapse imaging and single particle tracking.

### Electron microscopy

Detailed description for electron microscopy sample preparation (see **SI Materials and methods**, TEM and SEM sample preparation)

## Acknowledgements

We thank Prof. R. Sobotka and M. Foldynova (University of South Bohemia) for providing the anti-PilA1 antiserum; Prof. Masayuki Ohmori (Chuo University, Japan) and Dr. Shigeki Ehira (Tokyo Metropolitan University) for the generous gift of anti-adhA (cccS) antiserum. We were supported by the Carnegie Institution for Science, NSF (Grant MCB1331151) and Stanford University (Bio-X grant awarded to K. C. Huang, Bioengineering, and D.B.).

## Materials & Correspondence

Dr. Anchal Chandra

Prof. Devaki Bhaya

## References

1. Teschler JK, et al. (2015) Living in the matrix: assembly and control of Vibrio cholerae biofilms. Nat Rev Micro 13(5):255–268.

2. Guttenplan SB, Kearns DB (2013) Regulation of flagellar motility during biofilm formation. FEMS Microbiol Rev 37(6): 849–871.

3. Giltner CL, Nguyen Y, Burrows LL (2012) Type IV Pilin Proteins: Versatile Molecular Modules. Microbiol Mol Biol Rev 76(4):740–772.

4. Berry J-L, Pelicic V (2015) Exceptionally widespread nanomachines composed of type IV pilins: the prokaryotic Swiss Army knives. FEMS Microbiol Rev 39(1):1–21.

5. Mauriello EMF, Mignot T, Yang Z, Zusman DR (2010) Gliding Motility Revisited: How Do the Myxobacteria Move without Flagella? Microbiol Mol Biol Rev 74(2):229–249.

6. Burrows LL (2012) Pseudomonas aeruginosa Twitching Motility: Type IV Pili in Action. Annual Review of Microbiology 66(1):493–520.

7. López D, Vlamakis H, Kolter R (2010) Biofilms. Cold Spring Harb Perspect Biol 2(7). doi: 10.1101/cshperspect.a000398.

8. Bianca Brahamsha and Devaki Bhaya (2013) Motility in Unicellular and Filamentous Cyanobacteria. The cell biology of Cyanobacteria, Caister Academic Press, 233–262

9. Schuergers N, Wilde A (2015) Appendages of the Cyanobacterial Cell. Life 5(1):700–715.

10. Bhaya D, Bianco NR, Bryant D, Grossman A (2000) Type IV pilus biogenesis and motility in the cyanobacterium Synechocystis sp. PCC6803. Mol Microbiol 37(4):941–951.

11. Yoshihara S, et al. (2001) Mutational Analysis of Genes Involved in Pilus Structure, Motility and Transformation Competency in the Unicellular Motile Cyanobacterium Synechocystis sp. PCC6803. Plant Cell Physiol 42(1): 63–73.

12. Bhaya D, Watanabe N, Ogawa T, Grossman AR (1999) The role of an alternative sigma factor in motility and pilus formation in the cyanobacterium Synechocystis sp. strain PCC6803. Proc Natl Acad Sci USA 96(6):3188–3193.

13. Bhaya D, Takahashi A, Grossman AR (2001) Light regulation of type IV pilus-dependent motility by chemosensor-like elements in Synechocystis PCC6803. Proc Natl Acad Sci USA 98(13): 7540–7545.

14. Ikeuchi M, Ishizuka T (2008) Cyanobacteriochromes: a new superfamily of tetrapyrrole-binding photoreceptors in cyanobacteria. Photochem Photobiol Sci 7(10):1159.

15. Fiedler B, Borner T, Wilde A (2005) Phototaxis in the cyanobacterium Synechocystis sp. PCC 6803: role of different photoreceptors. Photochem Photobiol 81(6):1481–8.

16. Song J-Y, et al. (2011) Near-UV cyanobacteriochrome signaling system elicits negative phototaxis in the cyanobacterium Synechocystis sp. PCC 6803. Proc Natl Acad Sci USA 108(26):10780–10785.

17. Ursell T, Chau RMW, Wisen S, Bhaya D, Huang KC (2013) Motility Enhancement through Surface Modification Is Sufficient for Cyanobacterial Community Organization during Phototaxis. PLoS Comput Biol 9(9). doi: 10.1371/journal.pcbi.1003205.

18. Agostoni M, Montgomery BL (2014) Survival Strategies in the Aquatic and Terrestrial World: The Impact of Second Messengers on Cyanobacterial Processes. Life (Basel) 4(4):745–769.

19. Bhaya D, Nakasugi K, Fazeli F, Burriesci MS (2006) Phototaxis and Impaired Motility in Adenylyl Cyclase and Cyclase Receptor Protein Mutants of Synechocystis sp. Strain PCC 6803. J Bacteriol 188(20):7306–7310

20. Evans, ML., Chapman, MR. (2013). Curli Biogenesis: Order out of Disorder. Biochim Biophys Acta.10.1016/j.bbamcr.2013.09.010.

21. Busch A, Waksman G (2012) Chaperone-usher pathways: diversity and pilus assembly mechanism. Philos Trans R Soc Lond, B, Biol Sci 367(1592):1112–1122.

22. Fronzes R, Remaut H, Waksman G (2008) Architectures and biogenesis of non-flagellar protein appendages in Gram-negative bacteria. EMBO J 27(17):2271–2280.

23. Thanassi DG, Stathopoulos C, Karkal A, Li H (2005) Protein secretion in the absence of ATP: the autotransporter, two-partner secretion and chaperone/usher pathways of gram-negative bacteria (review). Mol Membr Biol 22(1–2):63–72.

24. Xu M, Su Z (2009) Computational prediction of cAMP receptor protein (CRP) binding sites in cyanobacterial genomes. BMC Genomics 10(1): 23.

25. Yoshimura H, Yanagisawa S, Kanehisa M, Ohmori M (2002) Screening for the target gene of cyanobacterial cAMP receptor protein SYCRP1. Mol Microbiol 43(4):843–853

26. Bhaya D, Takahashi A, Shahi P, Grossman AR (2001) Novel Motility Mutants of Synechocystis Strain PCC 6803 Generated by In Vitro Transposon Mutagenesis. J Bacteriol 183(20):6140–6143.

27. Geibel S, Waksman G (2014) The molecular dissection of the chaperone-usher pathway. Biochim Biophys Acta 1843(8):1559–1567.

28. Costa TRD, et al. (2015) Secretion systems in Gram-negative bacteria: structural and mechanistic insights. Nat Rev Micro 13(6):343–359.

29. Yoshimura H, et al. (2010) CccS and CccP are Involved in Construction of Cell Surface Components in the Cyanobacterium Synechocystis sp. strain PCC 6803. Plant Cell Physiol 51(7):1163–1172.

30. Nguyen, Y. et al. Pseudomonas aeruginosa Minor Pilins Prime Type IVa Pilus Assembly and Promote Surface Display of the PilY1 Adhesin. J Biol Chem 290, 601–611 (2015).

31. Ackermann M (2015) A functional perspective on phenotypic heterogeneity in microorganisms. Nat Rev Micro 13(8):497–508.

32. Jin F, Conrad JC, Gibiansky ML, Wong GCL (2011) Bacteria use type-IV pili to slingshot on surfaces. Proc Natl Acad Sci USA 108(31):12617–12622.

33. Anyan ME, et al. (2014) Type IV pili interactions promote intercellular association and moderate swarming of Pseudomonas aeruginosa. Proc Natl Acad Sci USA 111(50):18013–18018.

33. De La Fuente L, et al. (2007) Assessing Adhesion Forces of Type I and Type IV Pili of Xylella fastidiosa Bacteria by Use of a Microfluidic Flow Chamber. Appl Environ Microbiol 73(8):2690–2696.

34. Pesavento C, et al. (2008) Inverse regulatory coordination of motility and curli-mediated adhesion in Escherichia coli. Genes Dev 22(17):2434–2446.

35. Kaiser D, Crosby C (1983) Cell movement and its coordination in swarms of myxococcus xanthus. Cell Motility 3(3):227–245.

36. Fröls, S. et al. UV-inducible cellular aggregation of the hyperthermophilic archaeon Sulfolobus solfataricus is mediated by pili formation. Molecular Microbiology 70, 938–952 (2008).

37. Karatan, E. & Watnick, P. Signals, Regulatory Networks, and Materials That Build and Break Bacterial Biofilms. Microbiol Mol Biol Rev 73, 310–347 (2009).

38. Esquivel, R. N., Xu, R. & Pohlschroder, M. Novel Archaeal Adhesion Pilins with a Conserved N Terminus. J Bacteriol 195, 3808–3818 (2013).

